# QClus: A droplet-filtering algorithm for enhanced snRNA-seq data quality in challenging samples

**DOI:** 10.1101/2022.10.21.513315

**Authors:** Eloi Schmauch, Johannes Ojanen, Kyriakitsa Galani, Juho Jalkanen, Kristiina Harju, Maija Hollmén, Hannu Kokki, Jarmo Gunn, Jari Halonen, Juha Hartikainen, Tuomas Kiviniemi, Pasi Tavi, Minna U. Kaikkonen, Manolis Kellis, Suvi Linna-Kuosmanen

**Author notes:** To whom correspondence should be addressed: Suvi Linna-Kuosmanen. Email: <u>suvi.linna-</u>. These authors contributed equally to this work (Eloi Schmauch, Johannes Ojanen).

## Abstract

Single nuclei RNA sequencing (snRNA-seq) remains a challenge for many human tissues, as incomplete removal of background signal masks cell-type-specific signals and interferes with downstream analyses. Here, we present QClus, a droplet-filtering algorithm targeted toward challenging samples, using cardiac tissue as an example. QClus uses specific metrics such as cell-type-specific marker gene expression to cluster nuclei and filter empty and highly contaminated droplets, providing reliable cleaning of samples with varying number of nuclei and contamination levels. In a benchmarking analysis against seven alternative methods across six datasets consisting of 252 samples and over 1.9 million nuclei, QClus achieved the highest quality in the greatest number of samples over all evaluated quality metrics and recorded no processing failures, while robustly retaining numbers of nuclei within the expected range. QClus combines high quality, automation, and robustness with flexibility and user-adjustability, catering to diverse experimental needs and datasets.

## Introduction

Single-cell RNA sequencing (scRNA-seq) is a powerful tool for understanding the complex transcriptomes of heterogeneous cell populations ^1,2^. Single-nuclei RNA sequencing (snRNA-seq) uses the same principle but isolates nuclei instead of cells ^1^. Both methods are growing in use and popularity in biomedical research ^3^, however, snRNA-seq is particularly well suited for solid and frozen tissues, as well as trickier tissues, such as the human heart, where whole cells may be difficult to isolate and the use of single-cell approach essentially leads to biased cell yield due to the differences in cell sizes between the tissue cell populations (i.e., cardiomyocyte (CM) and non-cardiomyocyte (non-CM) populations in the heart) ^4,5^.

Droplet-based snRNA-seq works by encapsulating single nuclei in droplets, where each droplet contains RNA from one nucleus. However, ambient RNA contamination, such as cytoplasmic or cell-free RNA from the input solution, can contaminate the droplets. This is a significant concern for solid tissues, leading to misidentification of cell types and states ^6^. For example, in the human heart, CMs, the contractile units of the heart, are tightly bound together to facilitate their function ^7^, big in size, with high mitochondrial count and they produce high amounts of transcripts due to their size and metabolic activity ^9^, leading to high amount of cell debris and cytosolic RNA in the nuclei suspensions (**FIG 1A**), which can contaminate the snRNA droplets ^8,9^. This can result in mistaken interpretations of gene expression patterns ^6^, especially when CM transcripts are confused for genuine signals from other cell types. Such complications necessitate rigorous quality control in snRNA-seq workflows. In addition to heavily contaminated droplets, accurate exclusion of empty ones ^10^ is pivotal to avoid bias and ensure accuracy of subsequent analyses.

**Figure 1.**
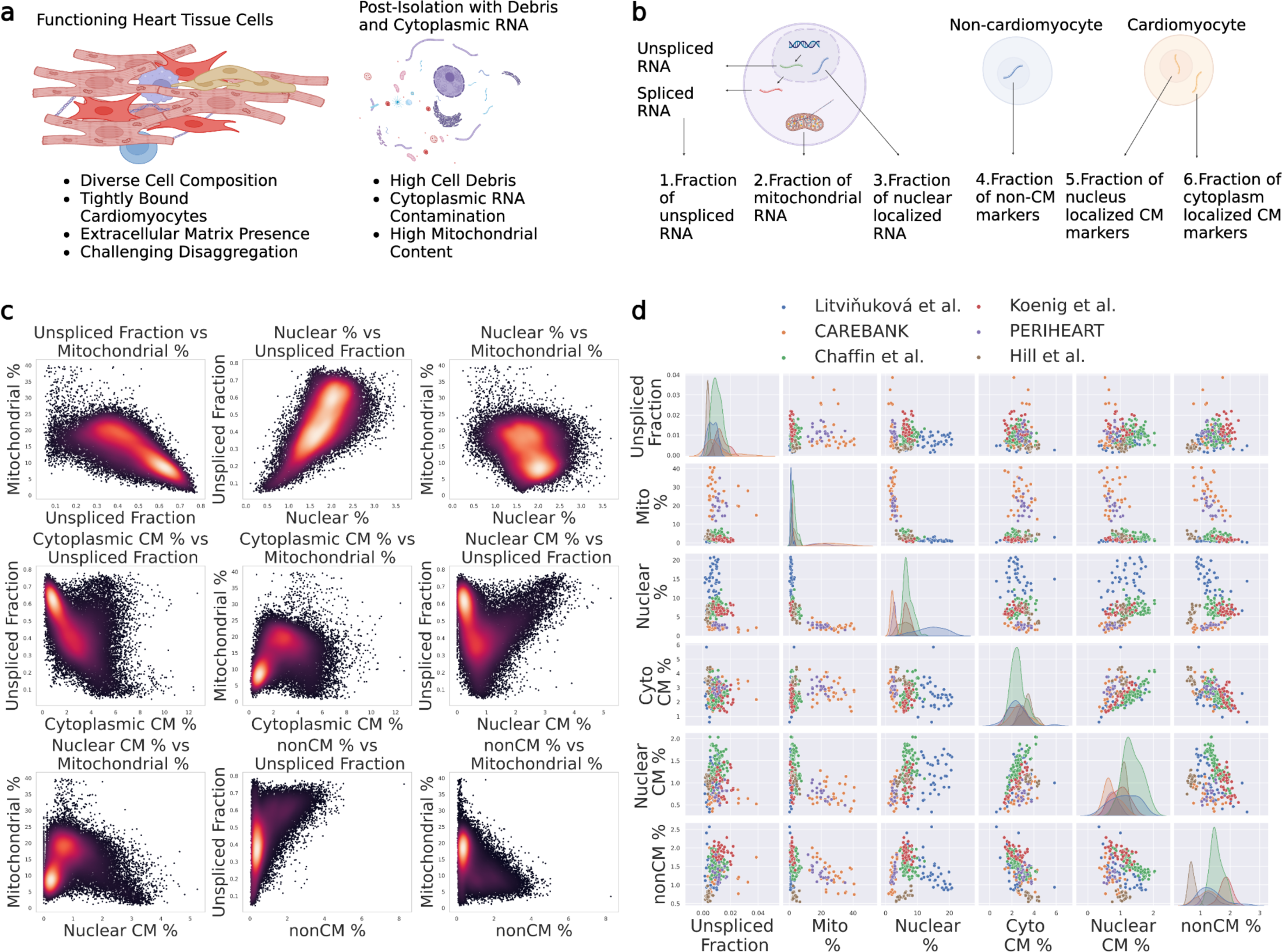
Use of general as well as cell type-specific quality metrics to model ambient RNA contamination in human heart snRNA-seq data. **a.** Illustration of tightly connected cardiomyocytes in living human cardiac tissue. Illustration of nuclei post attempted isolation, including significant amounts of debris and contamination. **b.** Illustration of general as well as cell type-specific metrics to model contamination that is heavily contributed to by cardiomyocytes. **c.** Plotting the metrics against each other for a single sample, colored by density, reveals areas of greater density in the feature space. **d.** Plotting the metrics across six datasets and 252 samples reveals high inter-sample variability.

Conventional methods for empty droplet removal in snRNA-seq often combine UMI distribution filtering (generally included in preprocessing programs, such as CellRanger) and mitochondrial fraction-based filtering ^6,10–14^. These techniques rely on clear distinction between empty and nuclei-containing droplets, which is typically true for samples with average contamination levels. However, in several tissue types, including the heart, elevated cytoplasmic contamination levels render standard removal methods inadequate ^6^. Recognizing this, recent years have seen the advent of bioinformatics tools aimed at identifying empty and highly contaminated droplets for removal or modeling contamination and correcting the expression count matrix data directly^6,11,15–17^.

In this study, we focused on determining the most effective method for automatic droplet filtering in extensive datasets of challenging samples with minimal manual input, using human cardiac tissue data as an example. After reviewing current methods ^6,8,11,15–18^ and testing them on the heart snRNA-seq data, we opted for a new approach to improve data quality by contamination-based nuclei filtering. Our primary objective was to develop a method that reliably and automatically works for larger datasets, even when such datasets demonstrate high inter-sample variability. We sought to construct a method that needed minimal manual input and no sample-level parameter tuning. To this end, we developed Quality Clustering (QClus), an innovative nucleus filtering method that employs unsupervised clustering of general as well as cell type-specific quality metrics. QClus, when compared to standard practices and prior algorithms, showed improved results in terms of quality metrics and robustness of nuclei retention across samples. QClus was specifically designed for nuclei calling instead of count decontamination, as our goal was to ensure the accurate identification of individual nuclei in the dataset, rather than attempting to cleanse or adjust the expression counts of detected transcripts. By focusing on the nuclei calling, we aimed to lay a foundation for subsequent analyses, ensuring that any interpretations or insights derived from the data would be based on correctly identified cellular components. By ensuring more dependable datasets, our methodology may help to offer deeper insights into human tissue biology.

## Results

### Description of Datasets

To understand how contamination is distributed in heart snRNA-seq data, we set out to explore the distribution of the known contamination metrics, such as mitochondrial fraction, as well as to develop novel metrics specific to the tissue type of interest (i.e., human heart tissue) that could be used during the quality control step of the workflow.

To explore the distribution of contamination metrics in human heart snRNA-seq datasets and benchmark droplet filtering methods, we collected 6 datasets. They consisted of four published datasets and two datasets of our own. The Chaffin *et al.* ^19^ data comprises 95 samples from the left ventricles of 11 patients with dilated cardiomyopathy, 15 with hypertrophic cardiomyopathy, and 16 control hearts. The Hill *et al.* ^20^ data provides 30 samples from nine pediatric patients with congenital heart disease (CHD) and four control hearts. The Koenig *et al.* ^21^ data includes 35 samples from the left ventricles of 17 patients with dilated cardiomyopathy and 28 control hearts. The Litviňuková *et al.* ^5^ data has 42 samples from six anatomical cardiac regions from 7 healthy hearts. Our own two datasets originate from a previous study, Linna-Kuosmanen *et al.* ^22^, including 50 right atrial appendage samples, from 15 patients with ischemic heart disease (IHD), 9 with myocardial infarction (MI), 11 with ischemic heart failure (IHF), 3 with non-ischemic heart failure, and 10 with valve disease.

### Description of Contamination Metrics

Two well-established universal snRNA-seq contamination metrics are the *unspliced* and *mitochondrial fractions* (**FIG 1B**). As the names suggest, the *unspliced fraction* measures the fraction of unspliced reads observed in the droplet and the *mitochondrial fraction* measures the fraction of reads originating from mitochondrial genes. Given the higher number of unspliced transcripts in the nucleus compared to cytoplasm, the metric tied to it is anticipated to show an inverse correlation with contamination, whereas *mitochondrial fraction* is expected to positively correlate with contamination, as mitochondria are only present in the cell cytoplasm. Both metrics have been established and utilized in previous research to measure ambient RNA levels ^6,10,16,17^. Expectedly, our findings in the heart data confirmed a strong negative correlation between the two metrics (**FIG 1C**), and the UMAP plots of the samples showed a central cluster exhibiting high levels of *mitochondrial* and low levels of *unspliced fraction* (**FIG S1**), corresponding to putative empty and highly contaminated droplets, thereby confirming the value of these metrics for measuring ambient RNA levels.

The third previously established metric in our study is the *nuclear fraction* (**FIG 1B**), which represents a nuclear-enriched gene expression fraction ^6,17^. *MALAT1* produces a transcript that localizes to nuclear speckles ^23^ and is believed to regulate the distribution and activity of splicing factors within these speckles. As nuclear-enriched genes possess a relative expression that should decrease in droplets with more cytoplasmic RNA, this fraction is expected to be inversely correlated with ambient RNA contamination in snRNA-seq. Consistently, we observed positive correlation with *unspliced fraction* and negative correlation with *mitochondrial fraction* for the metric (**FIG 1C**).

### Cardiomyocytes as a source of contamination

Cardiomyocytes, one of the most abundant cell types in cardiac tissue, possess a high RNA content in their cytoplasm due to their size and function. Accordingly, we observed CM-expressed genes to account for an important amount of the ambient RNA contamination in the samples (**FIG S2, FIG S3**). Thus, we hypothesized that some of these genes could be used as contamination metrics, provided that such a metric accounted for CM-nuclei-containing droplets, and not be biased against them.

To take this into account, we first needed a metric that would help us to distinguish CMs from non-CMs. Droplets that contain a single non-CM nucleus are expected to display high levels of gene expression corresponding to marker genes from one of the other major cell types. We thus defined a metric, *non-CM-specific gene expression fraction* (**FIG 1B**), for this purpose. It represents the fraction of reads aligning to the most highly expressed non-CM marker gene set (for details, including the definitions of gene sets, please see methods). After a closer look into the data, we observed that genes that were specific to other cell types than CMs, contributed to contamination at a much lower level than CMs (**FIG S3)**. A droplet that was enriched in one of the gene sets constituted by cell type marker genes was very likely to contain a nucleus of that cell type, instead of being empty. It was also less likely to be highly contaminated, since that would drive down this percentage, which was confirmed by the correlations with *unspliced fraction* and *mitochondrial fraction* in **FIG 1C**.

Droplets that contain a single CM nucleus are expected to display a high level of CM-specific gene expression. However, it can be difficult to distinguish true, high-quality CMs from contaminated droplets (either empty or containing a CM or non-CM nucleus) due to the high level of expression of CM-specific genes in the contamination. Interestingly, we observed CM-specific genes diverging into two groups: those contributing heavily to contamination and those contributing less (**FIG S4**), putatively corresponding to cytoplasm- and nuclear-enriched genes, respectively. Derived from this observation, we then defined two additional metrics: *cytoplasm-enriched CM-specific gene expression fraction* and *nuclear-enriched CM-specific gene expression fraction* (**FIG 1B**)*. Cytoplasm-enriched CM-specific gene expression fraction* can serve as a metric of contamination, as demonstrated by a strong correlation for the metric with *mitochondrial fraction* and negative correlation with *unspliced fraction* (**FIG 1C**). Conversely, *Nuclear-enriched CM-specific gene expression fraction* helped to distinguish true CM-containing droplets with low contamination from empty or highly contaminated droplets that may still contain significant amounts of CM marker gene expression due to the high contribution of CM-specific genes to ambient background RNA (**FIG 1C**).

### Contamination metric values vary significantly within and between dataset before preprocessing

After exploring and creating contamination metrics based on their specific distribution across droplets, we investigated their global distribution patterns across samples and datasets (**FIG 1D**) to establish an automated, universal filtering method. We observed a wide range of values for the created metrics within and across datasets, demonstrating the need for a method that automatically applies a flexible approach dependent on the sample-specific expression patterns and contamination levels. Full results for the mean values of the above quality metrics in unfiltered samples across the 252 samples can be found in **Table S1**.

### QClus efficiently removes empty droplets and highly contaminated nuclei

To answer the need for an automated method that can handle sample-specific expression patterns and contamination levels, we built QClus. QClus is centered on clustering droplets based on their defined quality metrics (**FIG 1C**). In addition, it uses CM-specific and non-CM-specific gene expression patterns as well as nuclear and cytoplasmic gene expression fractions in the metrics to distinguish CM nuclei, non-CM nuclei, highly contaminated droplets, and empty droplets, further improving the clustering. The QClus pipeline is divided into 5 steps:

***Step 1****: Input data*. The pipeline starts with data filtered by Cell Ranger (10X Genomics), which attempts to remove empty droplets using an algorithm based on the EmptyDrops ^11^ method (**FIG 2A**). Cell Ranger algorithm identifies low RNA content cells in samples with mixed cell populations. It initially uses a UMI count-based cutoff to detect high RNA content cells, followed by a detailed RNA profile analysis of remaining barcodes to separate actual cells from empty droplets. To illustrate this, we used a sample from the CAREBANK ^22^ dataset as an example.

**Figure 2.**
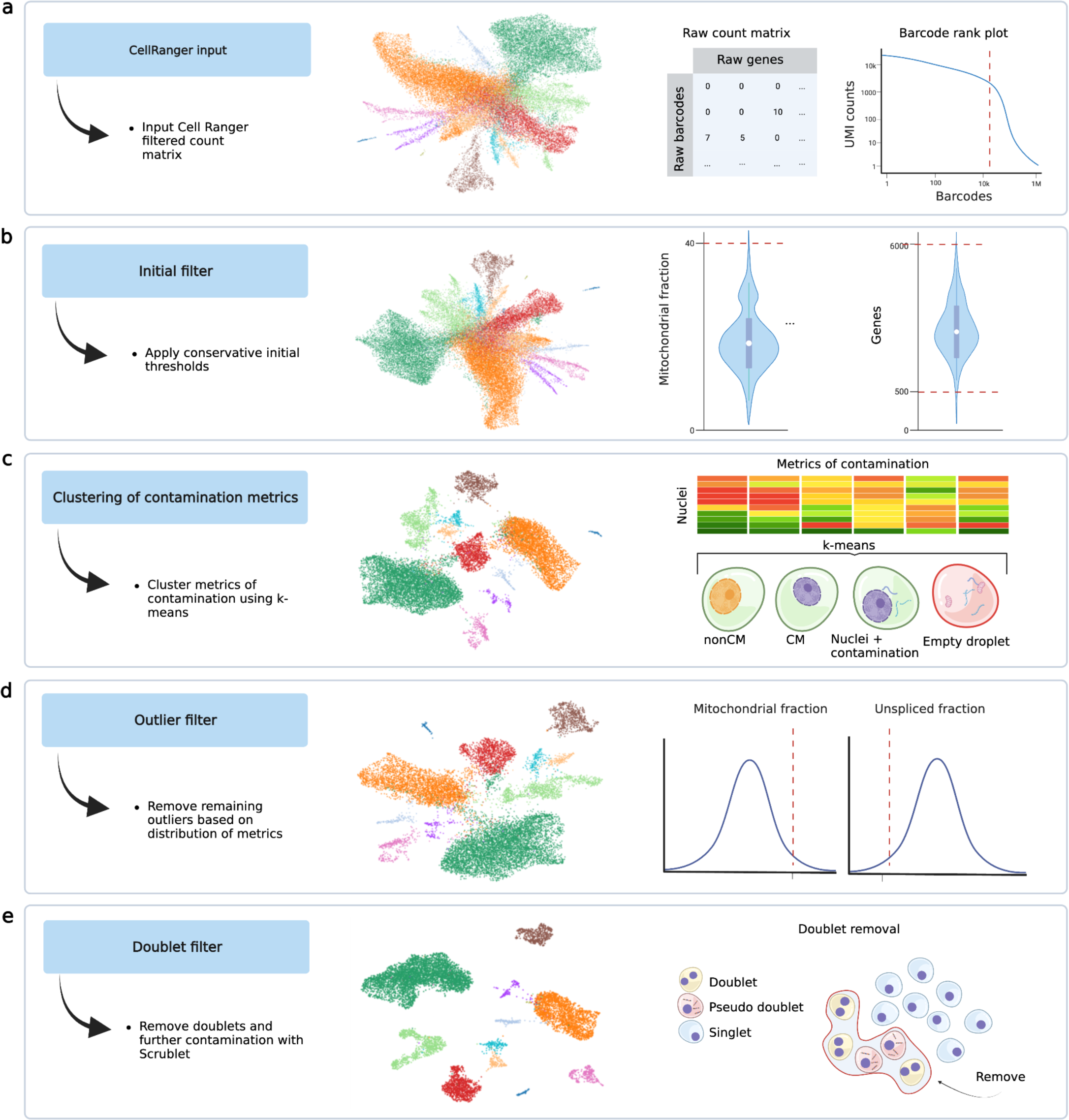
Overview of removal of empty droplets and highly contaminated droplets using QClus. **a.** The QClus algorithm begins with the pre-filtered count matrix that is output by Cell Ranger. **b.** A conservative initial filter is applied to remove tail ends of known quality metrics. **c.** General as well as cell type-specific quality metrics are calculated and clustered using k-means. The cluster displaying the lowest quality is removed by default. The user can choose to remove further clusters. **d.** An outlier filter based on the underlying distribution of mitochondrial percentage and unspliced fraction. **e**. Doublet filtering is applied to remove doublets as well as droplets exhibiting doublet-like expression due to the presence of a nonCM nucleus and CM derived ambient RNA. Created with BioRender.

After Cell Ranger filtering, the nuclei were found in a star shape on the dimension reduction map (UMAP), where all cell types seemed to originate from a common cluster. We hypothesized that the star center was composed of empty droplets and highly contaminated nuclei, as the cell types in the adult human heart are differentiated and therefore expected to form independent clusters. This hypothesis was confirmed by the high level of *mitochondrial fraction* and low level of *unspliced fraction* in the center of the star (**FIG S1**).

***Step 2****: Initial filtering.* The second filtering step removes droplets based on the detected number of genes and *mitochondrial fraction*, using high thresholds. The gene-level filtering ensures that all nuclei have enough genes to be of significant biological interest in the downstream analyses, whereas a mitochondrial threshold is used to remove clear outliers (**FIG 2B**).

***Step 3****: K-means clustering*. In this step, the method calculates the six contamination metrics defined earlier (**FIG 1B**; for details see methods). These values are then used as input for k-means clustering to identify four clusters (**FIG 2C**): **1)** non-CM nuclei with low levels of contamination, low CM-specific gene expression, both nuclear and cytoplasmic, and high non-CM-specific gene expression (**FIG S6**); **2)** CM nuclei with low levels of contamination, high CM-specific gene expression, both nuclear and cytoplasmic, and low non-CM-specific gene expression (**FIG S7**); and **3)** highly contaminated nuclei (**FIG S8**), and **4)** empty droplets (**FIG S9**, **FIG S10**), both of which have high contamination, high cytoplasmic CM-specific gene expression, low nuclear CM-specific gene expression, and low non-CM-specific gene expression. The default settings instruct QClus then to remove the empty droplets cluster only, but the user can choose to also remove the highly contaminated nuclei cluster, on a sample-by-sample basis (**FIG S10**). In most samples this step will remove the highest number of droplets and show the most significant improvements in sample quality.

***Step 4****: Outlier filtering*. The fourth filtering step removes highly contaminated nuclei in a more adjustable manner by identifying outliers based on the *unspliced* and *mitochondrial fraction* distributions within the sample (**FIG 2D**). In this process, outliers are selected and filtered based on a threshold determined by quantiles of the distribution within the non-CM cluster, defined as the cluster showing the highest level of the non-CM marker gene expression metric. This stage identifies and excludes those nuclei that deviate substantially from the expected distribution of the two metrics. It allows for a finer control over the quality and number of retained nuclei.

***Step 5****: Doublet filter*. To finalize the filtering, our pipeline uses the Scrublet algorithm^19^, which removes doublets. However, in addition to removing doublets, the Scrublet algorithm also removes highly contaminated nuclei in the heart, as remaining highly contaminated nuclei can appear as doublets (**FIG 2E**). These pseudo-doublets are non-CMs that have high levels of contamination (**FIG S11**) and will appear to contain RNA from CM cytoplasm (**FIG S12**).

After QClus preprocessing, the final UMAP showed a clear separation of the 11 major cell types observed in the illustrated sample (**FIG 2E**). No single metric alone predicted which nuclei were removed and which kept (**FIG S13**), suggesting that a multi-metric approach taken during the quality control maximizes the biological signal for downstream analyses.

### QClus outperforms other methods across multiple distinct quality metrics

To test our hypothesis regarding the effectiveness and versatility of the QClus method, we performed a comparative analysis of its performance against seven alternative droplet filtering methods across the six datasets. QClus was performed without the doublet filtering step to provide a fair comparison against other methods which do not include doublet filtering. In addition, QClus was set to its default mode regarding ***step 3***, removing only the empty droplet cluster in this step (**FIG S6**), allowing for fully automated execution. Utilizing the six datasets, we selected four distinct quality metrics for evaluation: *unspliced fraction*, *mitochondrial percentage*, total counts, and the number of genes expressed (for full results, see **Table S2**). For each sample-method combination, the metrics were normalized on a per-sample basis (for full scaled results, see **Table S3**). This normalization ensured the standardized comparison of the performance of each method, across all samples.

Across all four QC metrics, QClus displayed the highest number of samples with the best result for that metric (**FIG 3A**), outperforming other methods in 138 out of 252 samples (54.76%) for *unspliced fraction*, in 116 samples (46.03%) for *mitochondrial percentage*, 78 samples (30.95%) for total counts, and 96 samples (38.10%) for the mean number of genes expressed per nuclei.

**Figure 3.**
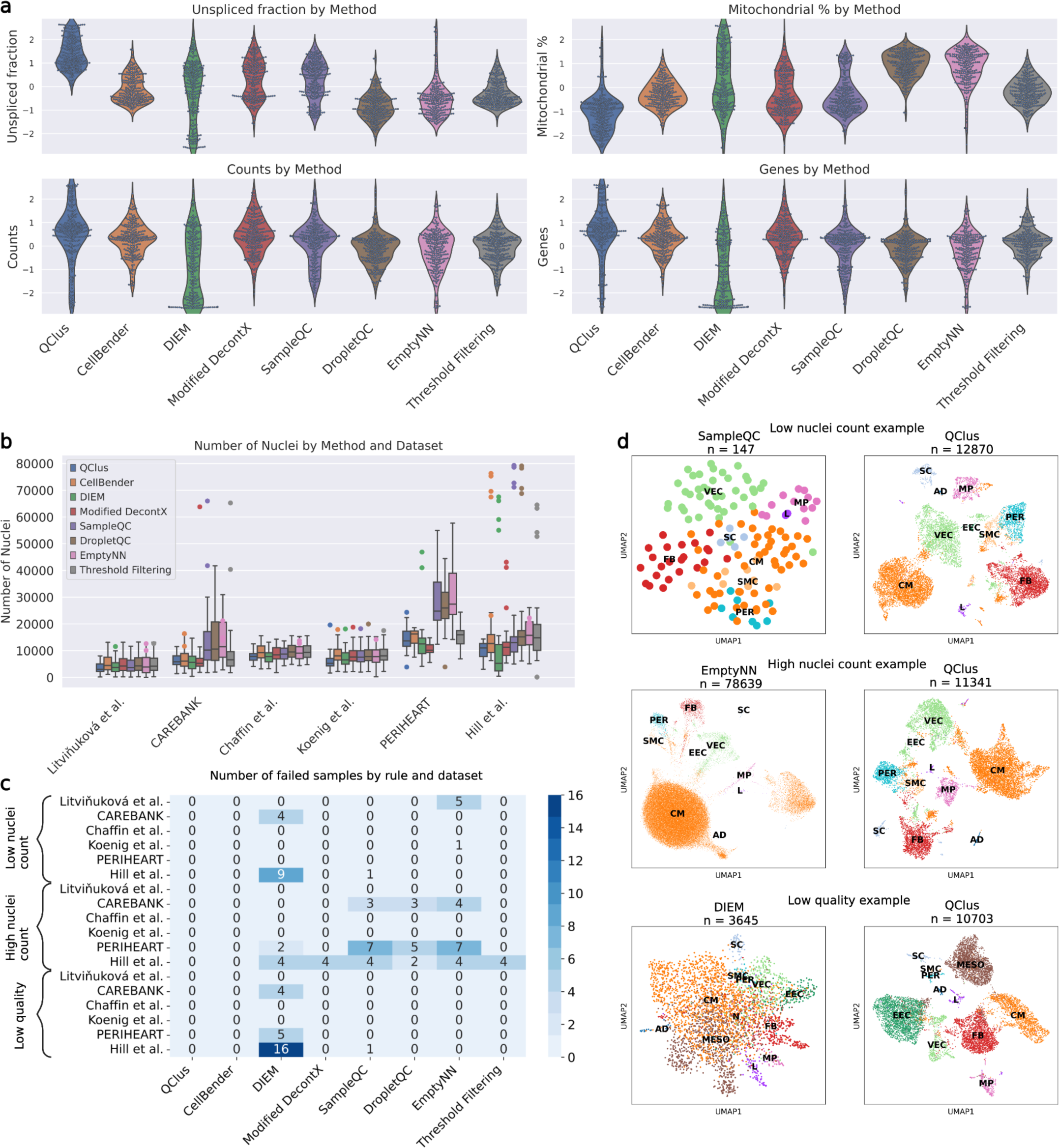
Benchmarking QClus against alternative methods shows improved quality metrics and robust nuclei retention. **a.** Comparing achieved quality of six published methods as well as a control method of traditional quality metric threshold filtering demonstrates improved quality in most samples. **b.** QClus retains a comparable number of nuclei compared to some methods, while some others retain amounts of nuclei significantly outside the range of expectations. **c.** All methods outside of QClus and CellBender report failed sample processing for some samples. **d.** Three examples of failed samples by alternative methods compared against the same samples processed with QClus. Examples from Hill et al. (Hi_WU198RV_rep1), Hill et al. (Hi_WU13235_rep1), and CAREBANK (CB-Q34), respectively.

Another important metric in the assessment of droplet filtering methods is the resulting number of kept nuclei. Due to the distribution of quality metrics across nuclei, a more stringent threshold decreases the number of nuclei and increases their quality. There is thus an inherent trade-off between the number of nuclei retained and the quality achieved. While retaining more nuclei can potentially enhance the richness of the dataset, it might do so at the expense of lower quality, which could lead to noise or false conclusions. Conversely, being overly stringent might result in the omission of high-quality nuclei, thereby limiting the scope of insights that could be drawn. Investigation of this aspect across the datasets and methods revealed that QClus, although retaining fewer nuclei compared to some methods, such as EmptyNN ^15^, DropletQC ^17^, and SampleQC ^16^, remained within the expected range for nuclei retention across various datasets (**FIG 3B**). In addition, compared to other methods, the nuclei retention counts appeared more stable. Given the consistent nature of experimental procedures within each dataset, large fluctuations of nuclei retention could be indicative of a less reliable filtering.

Unlike some of the methods, the adaptability of the QClus algorithm allows its parameters to be adjusted to fit specific research needs, whether the user wishes to retain more nuclei or take a more stringent approach. This offers a flexible balance between nuclei retention and quality, catering to various experimental and analysis requirements.

To further benchmark and compare the eight preprocessing methods, we established criteria that, when fulfilled, would indicate a processing failure—i.e., an unacceptable result for the respective sample. The method-wise numbers on how many samples failed for each rule are shown in **FIG 3C** and **Table S4**. The first criteria considered, was a ‘low nuclei count’, which red-flagged cases where a considerably greater number of nuclei of comparable quality could have been retained, i.e., any method that retained fewer than 3,000 nuclei for a particular sample, while another method was able to retain over four times as many nuclei with an unspliced fraction within 10% for the same sample. Twenty sample-method cases fell into this category. These failures were spread across DIEM ^6^ (13 samples), EmptyNN ^15^ (6 samples), and SampleQC ^16^ (1 sample). An example from the Hill *et al.* ^20^ dataset is shown in **FIG 3G**, where SampleQC retained 147 nuclei, while QClus was able to retain 12,870 nuclei of good quality (another example is shown in **FIG S14)**.

The second criteria considered, was a ‘high nuclei count’, which flagged samples that retained more than 30,000 nuclei after preprocessing. Given the experimental design of the datasets, parameters of the preprocessing methods (such as the *expected cells* parameter of CellBender ^24^) used in the publications, and observed ranges of retained nuclei in the original publications, samples were expected to contain between 3,000 and 10,000 nuclei. Of the analyzed sample-method combinations, 53 met this condition, signaling a failure in quality control. These failures were distributed across six methods: Modified DecontX ^18^ (4 samples), DIEM ^6^ (6 samples), DropletQC ^17^ (10 samples), EmptyNN ^15^ (15 samples), traditional preprocessing (4 samples), and SampleQC ^16^ (14 samples). An example from the Hill *et al.* ^20^ dataset is shown in **FIG 3G**, where EmptyNN retained 78,639 nuclei, while QClus retained 11,341 nuclei (another example is shown in **FIG S14)**.

The third criteria used, was a ‘low quality’. A method was considered to have failed, when the median unspliced fraction for the retained nuclei of a method-sample combination was less than 0.3, and another method retained a greater number of nuclei from the same sample with a median unspliced fraction of at least 0.2 higher. Thus, the same number of nuclei of significantly higher quality could be identified using an alternative method. Twenty-six sample-method combinations failed using this criteria, namely DIEM ^6^ (25 samples) and SampleQC ^16^ (1 sample). An example from the CAREBANK dataset is shown in **Figure 3G**, where DIEM retained 3,645 nuclei with a mean unspliced fraction of 0.22 and no clear cell type separation on the UMAP, while QClus retained 10,703 nuclei with an unspliced fraction of 0.64 and clear separation (another example is shown in **FIG S14)**.

Taken together, QClus outperformed other methods by displaying the highest number of samples with the best results in quality while remaining within the expected range for nuclei retention across various datasets, with more stable nuclear retention counts compared to other methods. Based on the criteria for processing failures, only two of the methods, QClus and CellBender ^24^, exhibited no failures across any of the six datasets. However, while CellBender processed samples passed the set failure criteria, there were several samples in which it did not perform a good quality filtering. These same samples were found to be processed better with QClus filtering. In an example shown in **Figure 4A**, CellBender filtering resulted in low cell type separation and distribution, whereas after QClus, cell types were found in higher abundance and balanced composition (**FIG 4B**).

**Figure 4.**
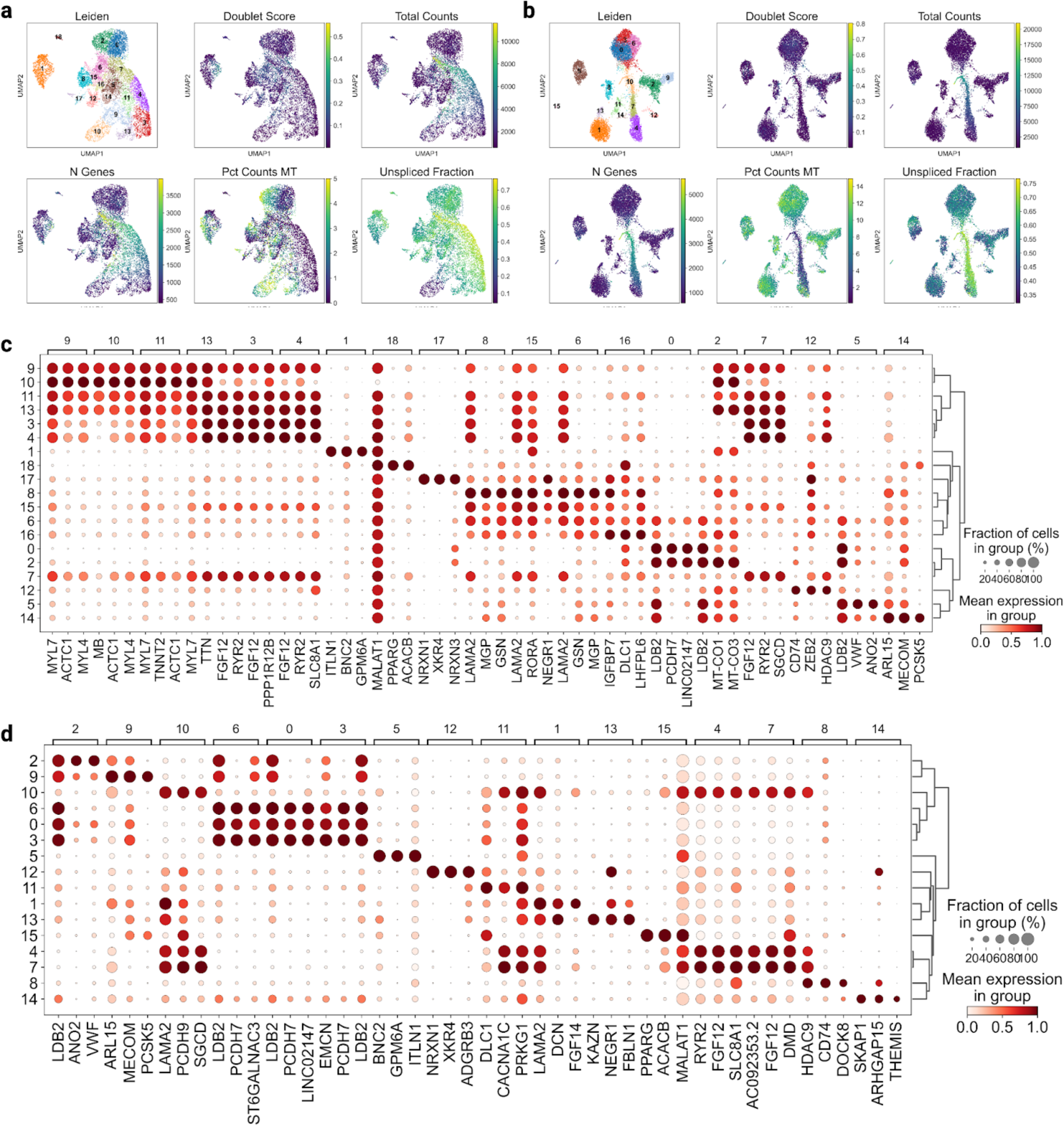
Comparison of CellBender and QClus. **a.** UMAPs and dotplot for CellBender processing of a sample of the PERIHEART dataset, representing samples where CellBender failed to achieve satisfactory quality. **b.** UMAPs and dotplot of QClus processing of the same sample, representing samples where significantly improved separation and composition of cell types was seen with the QClus processing. **c-d.** Dot plot of top three genes per leiden group for CellBender (**c**) and QClus (**d**) processing of the same sample.

## Discussion

QClus is a droplet filtering method that is tailored to overcome challenges in snRNA-seq processing tissue-specifically. The benchmarking results advocate for specialized preprocessing, especially when grappling with samples that contain good-quality nuclei covered with substantial contamination.

The strength of QClus lies in its ability to process heterogeneous datasets in an automated way while adapting to variations in-sample characteristics, such as nuclei number, overall quality, and contamination levels. The adaptability reduces the need for meticulous manual oversight required in other similar methods, and our benchmarking attests to the performance of QClus in unsupervised scenarios. As the method is designed to work tissue-specifically, we focused here on one specific tissue type known to be challenging, the human heart tissue and geared our approach to heart-specific features that our research corroborated. However, the method is adaptable to other tissue types by modifying the list of marker genes or the input metrics. The flexibility of the method allows the parameters of the algorithm to be tweaked globally or at the level of individual samples, providing researchers with the opportunity to balance between nuclei retention and contamination control, based on the specific biological questions at hand.

As QClus is specifically designed to function without supervision and tailored for high-throughput datasets, where manual intervention is not feasible, it is plausible that some benchmarked methods would have shown enhanced performance had they been given more manual adjustments tailored to each sample. However, such a procedure would deviate from our primary objective, which was to identify a method that reliably and automatically works for extensive datasets with minimal manual input in a specific tissue context. With the advent of novel techniques, the decreasing cost of sequencing, and the generation of larger datasets, there is an unmistakable demand for fully automated methods that can efficiently handle large volumes of data while maintaining accuracy and reliability.

QClus, though effective, has room for growth. Accounting for contamination distributed within cell type populations might improve accuracy, given that we observe varying levels even within non-CMs. Integrating QClus with a decontamination algorithm, such as CellBender ^24^ or DecontX ^18^, might further enhance its efficiency, establishing an integrated solution that both filters and cleans up the retained nuclei. Thus, the optimal strategy could involve a careful and balanced approach to both nuclei calling and decontamination. The specific choice and combination of methods and strategies might be guided by the tissue type, cellular complexity, and specific questions asked in a study.

In conclusion, challenging tissues, exemplified by cardiac tissue in our study, highlight the need for tissue-specific preprocessing methodologies that can handle large datasets containing samples of varying qualities in an automated fashion. The idiosyncrasies of contamination across different tissues and even disorders demand tailored solutions. QClus emerges as a potent tool for such preprocessing – as shown in cardiac tissue, with promising avenues for adaptation to other tissue types – fostering an era of more precise, tailored snRNA-seq analyses.

## Methods

### Datasets and preprocessing

For the 4 external dataset, the FASTQ files from each sample were downloaded from their respective repository. The Hill *et al.* ^20^ and Koenig *et al.* ^21^ data were obtained from GEO (respectively GSE203275 and GSE183852). The Litviňuková *et al*. ^5^ dataset was obtained from ENA (https://www.ebi.ac.uk/ena/browser/view/PRJEB39602). The Chaffin *et al.* ^19^ data was obtained from dbGAP (https://www.ncbi.nlm.nih.gov/projects/gap/cgi-bin/study.cgi?study_id=phs001539.v1.p1) with proper authorizations.

More information about the datasets, their experimental processing, the patient’s conditions, can be found in their respective papers ^5,19–21^.

All samples from each dataset were processed in parallel using a dedicated high-performance computing (HPC) cluster in our laboratory, managed via the SLURM workload manager.

Package versions were as followed:

- CellBender 0.2.2
- Velocyto 0.17.17
- Scanpy 1.9.2
- SampleQC 0.6.6
- Diem 2.4.1
- Celda 1.14.1 for the use of DecontX
- DropletQC 0.0.0.9000
- EmptyNN 1.0

CellRanger v7 was run for all samples, with default parameters, and the GRCh38 reference file provided. Then, velocyto ^25^ was run using the *run10x*-command, providing as input, in addition to the CellRanger output, the genome annotation file from the CellRanger GRCh38 reference genome. In addition, as advised by the velocyto ^25^ tutorial, the repeat regions were masked and a repeat masker file was downloaded from UCSC genome browser, and provided as input.

To benchmark the effectiveness of QClus against six previously published methods of nuclei filtering, in addition to an adapted implementation of a decontamination method, each method was run for all samples through a custom python pipeline, which outputted the list of barcodes to keep. All methods, including QClus, were executed with their default parameters whenever possible, following available tutorials in their respective documentation. Specific input was made only when required or to ensure consistency across comparisons. Details about specific parameter settings for each method are given below. As a base filtering, CellRanger default filtering (whenever a method did not specifically call for the unfiltered count matrix as input), in addition to conservative thresholds of commonly used QC metrics (total counts, total genes, and *mitochondrial fraction*), was applied. This ensures applicability of the benchmark results. Exact values for these thresholds are given in **Table S5**.

### QClus

#### Initial filter

To remove droplets with poor QC metrics, the lower bound of the number of detected genes is set at 500, the upper bound of the number of detected genes at 6000, and the initial cutoff for *mitochondrial fraction* to be 40%.

#### Clustering features: Unspliced fraction

After annotating reads as “spliced”, “ambiguous”, or “unspliced” using Velocyto ^25^, a “fraction_unspliced” metric is calculated for each droplet. It is the total number of unspliced reads over the total number of reads (spliced, unspliced and ambiguous) in each droplet.

#### Clustering features: *Mitochondrial fraction*

This metric corresponds to the fraction of reads aligning to the mitochondrial genome (*MT-ND1, MT-ND2, MT-CO1, MT-CO2, MT-ATP8, MT-ATP6, MT-CO3, MT-ND3, MT-ND4L, MT-ND4, MT-ND5, MT-ND6, MT-CYB*), for each droplet. The metric is calculated using the Scanpy ^26^ function calculate_qc_metrics() on raw counts.

#### Clustering features: *Nuclear fraction*

This metric corresponds to the fraction of reads aligning to the following genes: *MALAT1, NEAT1, FTX, FOXP1, RBMS3, ZBTB20, LRMDA, PBX1, ITPR2, AUTS2, TTC28, BNC2, EXOC4, RORA, PRKG1, ARID1B, PARD3B, GPHN, N4BP2L2, PKHD1L1, EXOC6B, FBXL7, MED13L, TBC1D5, IMMP2L, SYNE1, RERE, MBD5, EXT1, WWOX*. These were chosen as they exhibit high correlation with *MALAT1*, which is known to be highly expressed in the nucleus. The metric is calculated using the Scanpy function calculate_qc_metrics() on logarithmized counts. Logarithmization is performed using the Scanpy function log1p().

#### Clustering features: Nuclear CM marker genes

A set of genes found to be specifically expressed in CMs, but absent from ambient RNA contamination were selected as highly expressed in CM nuclei: *RBM20, TECRL, MLIP, CHRM2, TRDN, PALLD, SGCD, CMYA5, MYOM2, TBX5, ESRRG, LINC02248, KCNJ3, TACC2, CORIN, DPY19L2, WNK2, MITF, OBSCN, FHOD3, MYLK3, DAPK2, NEXN*. Droplets with a high level of expression of these genes are expected to contain CM nuclei. The metric is calculated using the Scanpy function calculate_qc_metrics() on raw counts.

#### Clustering features: Cytoplasmic CM marker genes

A set of genes found to be specifically expressed in CMs, but present in high level from ambient RNA contamination were selected as highly expressed in CM cytoplasm: *TTN, RYR2, PAM, TNNT2, RABGAP1L, PDLIM5, MYL7, MYH6*. The metric is calculated using the Scanpy function calculate_qc_metrics() on raw counts.

#### Clustering features: Cell type specific fractions

For each of the eleven remaining cell types, a set of genes was selected as markers of these cell types from wilcoxon rank sum test differential expression analysis (**Table S5**). The fraction of reads aligning to those sets of genes was calculated for each droplet,using the Scanpy function calculate_qc_metrics() on raw counts. Next, for each droplet, the maximum value out of those 11 metrics was selected.

#### Clustering method

Once the clustering features have been computed for the remaining nuclei, the features are scaled using the MinMaxScaler() function implemented in the scikit-learn ^27^ package. Next, the k-means algorithm was used, as implemented in the scikit-learn package, to partition the nuclei into four clusters, using all 6 clustering features. By default, the cluster demonstrating the lowest mean value of *unspliced fraction* is removed, as this cluster is predicted to contain empty droplets. However, the user can choose to remove any number of clusters. The default number of clusters is 4, but that number can be changed to adapt the software to different contamination profiles (**FIG S10**).

#### Outlier filtering

Filtering threshold values are calculated for the fraction of unspliced reads in addition to *mitochondrial fraction*, based on the distribution of these metrics in the r non-CM cluster, defined as the cluster with the highest mean value for the pct_counts_nonCM metric. The unspliced fraction threshold is chosen as the lower quartile minus 0.1. The *mitochondrial fraction* threshold is chosen as the upper quartile plus 0.05. These values can be adjusted by the user. Droplets are then filtered out if they go beyond these thresholds in both metrics (below the threshold for *unspliced fraction* and above for *mitochondrial fraction*).

#### Doublet removal

The scrublet algorithm is used in each sample, and droplets whose doublet score exceeds 0.1 are removed. This filters out both doublets and highly contaminated droplets, whose distribution can resemble doublets in terms (**FIG S12**).

### Other Methods

#### Traditional Threshold Filtering Based on QC Metrics Only

As a baseline, we employed standard QC filtering (threshold filtering) that is routinely run in snRNA analysis, and often not accompanied by more filtering methods. . Metrics used were *mitochondrial fraction*, *total counts*, *total genes*. The exact threshold values utilized were set based on the details given in the respective studies and can be reviewed in **Table S6**.

#### DIEM

DIEM ^6^ is a droplet filtering method that is based on a model of RNA expression, taking into account contamination and cell type specific patterns, using a multinomial distribution. The initialization phase requires users to set a stringent count threshold based on a barcode rank plot, subsequently tagging nuclei below this threshold as debris, meaning they are expected to represent the profile of ambient RNA contamination. Using the expectation-maximization algorithm, DIEM determines model parameters. Each droplet receives a score reflecting its expression of genes attributed to the debris set, enabling users to implement further filtration based on the debris score. The debris score was calculated using default settings and a debris score cut-off of 0.5, which is the default value, was set to filter out empty and highly contaminated droplets.

#### Modified DecontX

DecontX ^18^ employs a Bayesian framework to decipher the distribution of background contamination and ascertain contamination levels in each droplet. This model is then used to remove contamination from the gene expression matrix. In this study, we focus on droplet filtering and not decontamination *per se*. Thus, the contamination score derived from DecontX results was used to filter droplets. Droplets that had a contamination score of more than 0.43 were removed. The threshold was established to compare DecontX results to QClus results, as it resulted in the same number of droplets being filtered out in the PERIHEART ^22^ dataset.

#### EmptyNN

EmptyNN ^15^ is a cell-calling algorithm to differentiate between droplets that contain cells and those that are empty or cell-free. It utilizes a PU (Positive and Unlabeled) learning bagging strategy, based on the idea that barcodes with very low total counts are likely to represent genuine cell-free droplets. It has also been benchmarked with snRNA-seq data.

#### SampleQC

SampleQC ^16^ fits a Gaussian mixture model spanning multiple samples, subsequently filtering outlier nuclei. The metrics we used as inputs were counts, genes, mitochondrial fraction, and unspliced fraction. The ‘k_all’ argument was set to ‘k_all=2’, which was observed to fit the data well in most cases. The method suggests testing different values of that parameter between groups of samples. However, further optimization is outside of the scope of our study, as it renders benchmarking more subjective: we thus concentrated on fully unsupervised methods, and setting parameters to their default value.

#### DropletQC

DropletQC ^17^ utilizes both splicing fraction and total detected UMIs to discern among empty droplets, intact nuclei, and contaminated nuclei. Estimated thresholds are employed to identify nuclei positioned above or below these designated values. We executed the identify_empty_drops function of the package to identify and remove empty droplets.

#### CellBender

CellBender ^24^ is a tool designed to decontaminate data by constructing a probabilistic model to differentiate between true biological counts and background noise in the observed feature count matrix, using a deep generative model. CellBender was run with the expected-cells parameter set to 5000, the total-droplets-included parameter set to 40000, the *fpr* parameter set to 0.01, and the epochs parameter set to 150. CellBender uses its modeling of contamination to identify empty and non-empty droplets, in addition to the decontamination itself. This droplet selection feature was used for the benchmarking. However, the contamination removal part of the output was not used, and the original count matrix is used to calculate quality metrics for benchmarking.

### Benchmarking

After running each of the droplet filtering methods on all samples, for all datasets, quality metrics were calculated at the sample-level. Standard scaling was performed with StandardScaler from scikit-learn to standardize quality metrics for each unique sample. Each sample data was isolated and scaled independently.Embeddings for each sample-method combination were constructed using Scanpy, using standard dimension reduction and embedding procedures, as laid out in the Scanpy tutorial. The sc.pp.filter_genes function filtered genes in the AnnData object present in fewer than 10 nuclei. Subsequently, highly variable genes were identified using the sc.pp.highly_variable_genes function with parameters min_mean=0.005, max_mean=5, and min_disp=0.5. The dataset was then narrowed down to these highly variable genes.Data corrections for total counts and the percentage of mitochondrial counts were applied with sc.pp.regress_out. The sc.pp.scale function scaled the data to a maximum value of 10.

## Code availability

Qclus is a ready-to-use python package, which is suited for integration with Scanpy processed single nuclei data, taking the AnnData object as input. Instructions to download and install the package in addition to its source code can be found on github: https://github.com/scHEARTGROUP/qclus.

## Acknowledgements

This work was supported by the Aarne Koskelo Foundation [to E.S. and S.L.K.]; the Academy of Finland [grant number 342074 to S.L.K.]; the Antti and Tyyne Soininen Foundation [to E.S.]; the Doctoral Program of Molecular Medicine, University of Eastern Finland [to E.S.]; the Finnish Foundation for Cardiovascular Research [to M.K. and S.L.K.]; the Finnish Academy of Science and Letters [to E.S.]; the Maud Kuistila Memorial Foundation [to S.L.K.]; the Orion Research Foundation [to E.S and S.L.K.]; the Saastamoinen Foundation [to E.S.]; the Sigrid Juselius Foundation [to M.U.K. and S.L.K.]; and the Yrjö Jahnsson Foundation [to E.S. and S.L.K.].

For the Chaffin *et al.* dataset, we thank the Broad Institute for generating high-quality sequence data with Patrick Ellinor as the PI. The dataset used in this manuscript (referenced as the Chaffin *et al.* dataset) was obtained from dbGaP at http://www.ncbi.nlm.nih.gov/gap through dbGaP accession number phs001539.

## Notes

### Competing Interest Statement

The authors have declared no competing interest.

### Summary of Updates

The revised manuscript enhances the presentation of the QClus method. This update introduces a more thorough benchmarking analysis that compares QClus' performances with seven alternative droplet-filtering algorithms. The analysis spans six single nuclei RNA-seq heart datasets, totaling 252 samples and over 1.9 million nuclei. The manuscript's revisions include clarified figures and refined methodological descriptions for better comprehension and reproducibility. Additionally, a GitHub repository link has been provided, permitting to install and use the QClus Python package.

