## Supplementary Figures for "QClus: A droplet-filtering algorithm for enhanced snRNA-seq data quality in challenging samples"

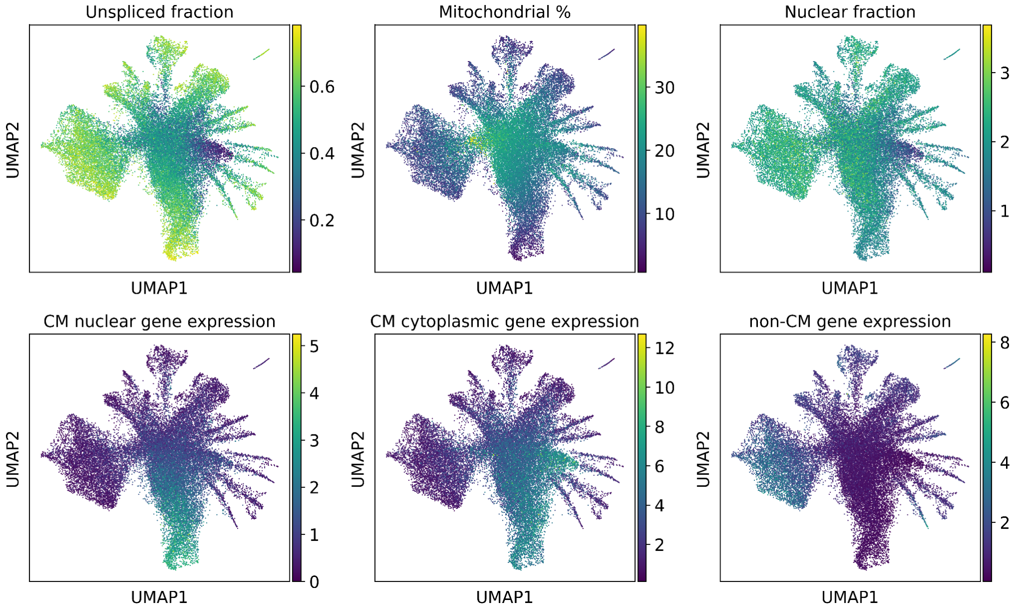


**Figure. S1**. Distribution of the metrics that are used by QClus across an unfiltered sample (CB-S00) from our previous study (doi: https://doi.org/10.1101/2021.06.23.449672).

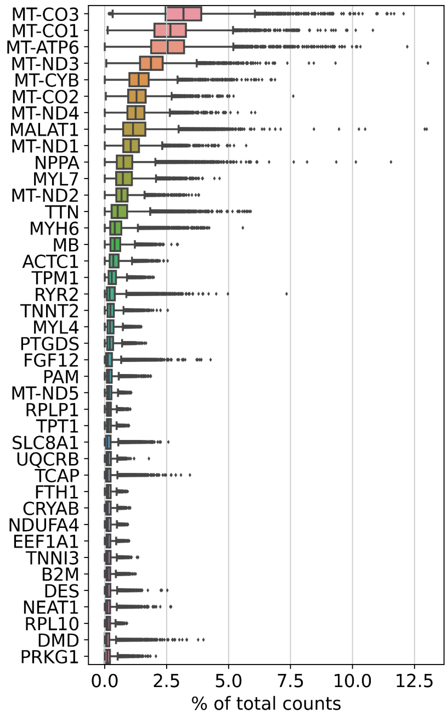


**Figure. S2**. Most abundant genes in empty droplets-cluster. A high presence of reads stemming from mitochondria and cardiomyocyte marker genes is observed.

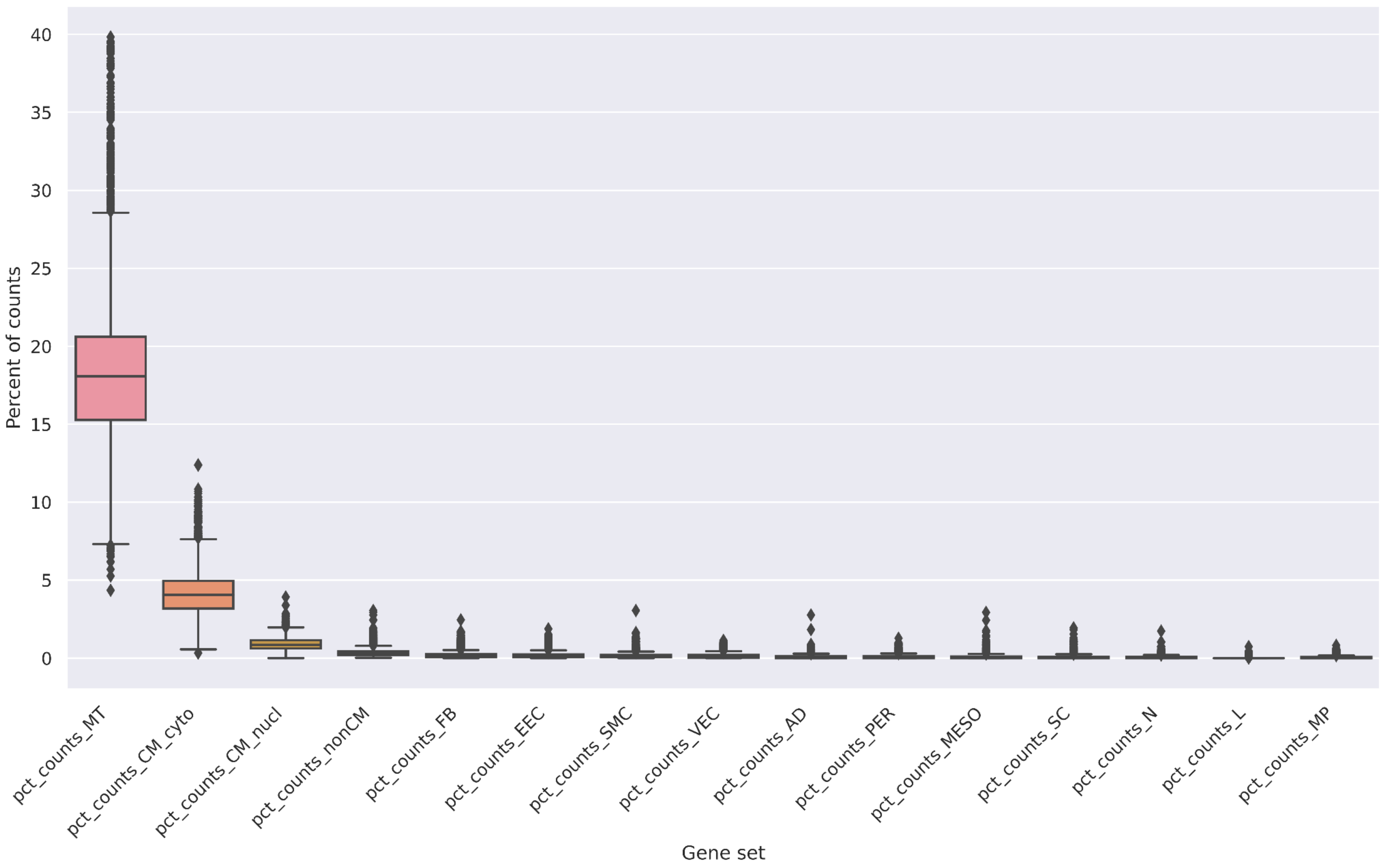


**Figure. S3**. Fraction of reads aligning to the identified gene sets in empty droplets. A high fraction of reads aligns to either mitochondrial genes or genes identified as cytoplasm-enriched cardiomyocyte genes.

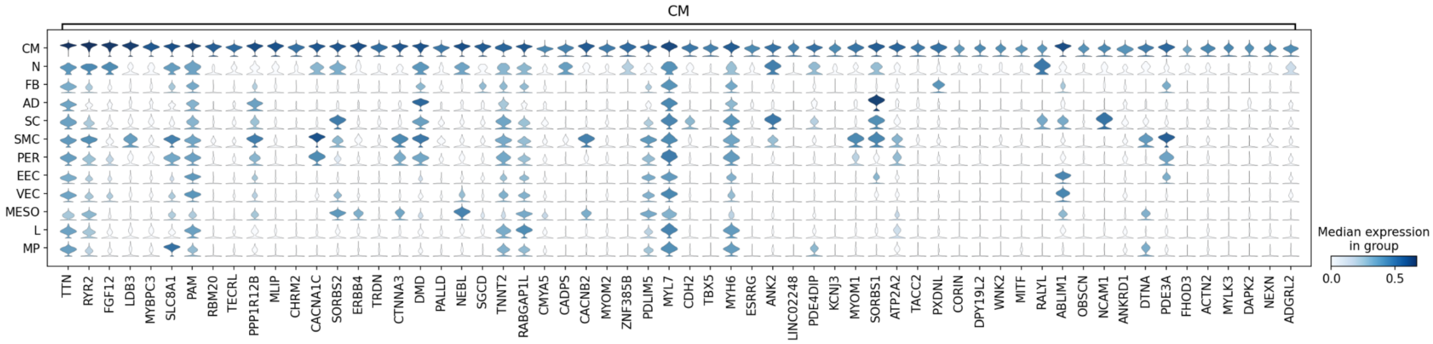


**Figure. S4**. Cardiomyocyte-specific gene expression across cell type populations. Cardiomyocyte-specific genes can be observed to be either highly specific to only cardiomyocytes or also present to a significant degree in other cell types.

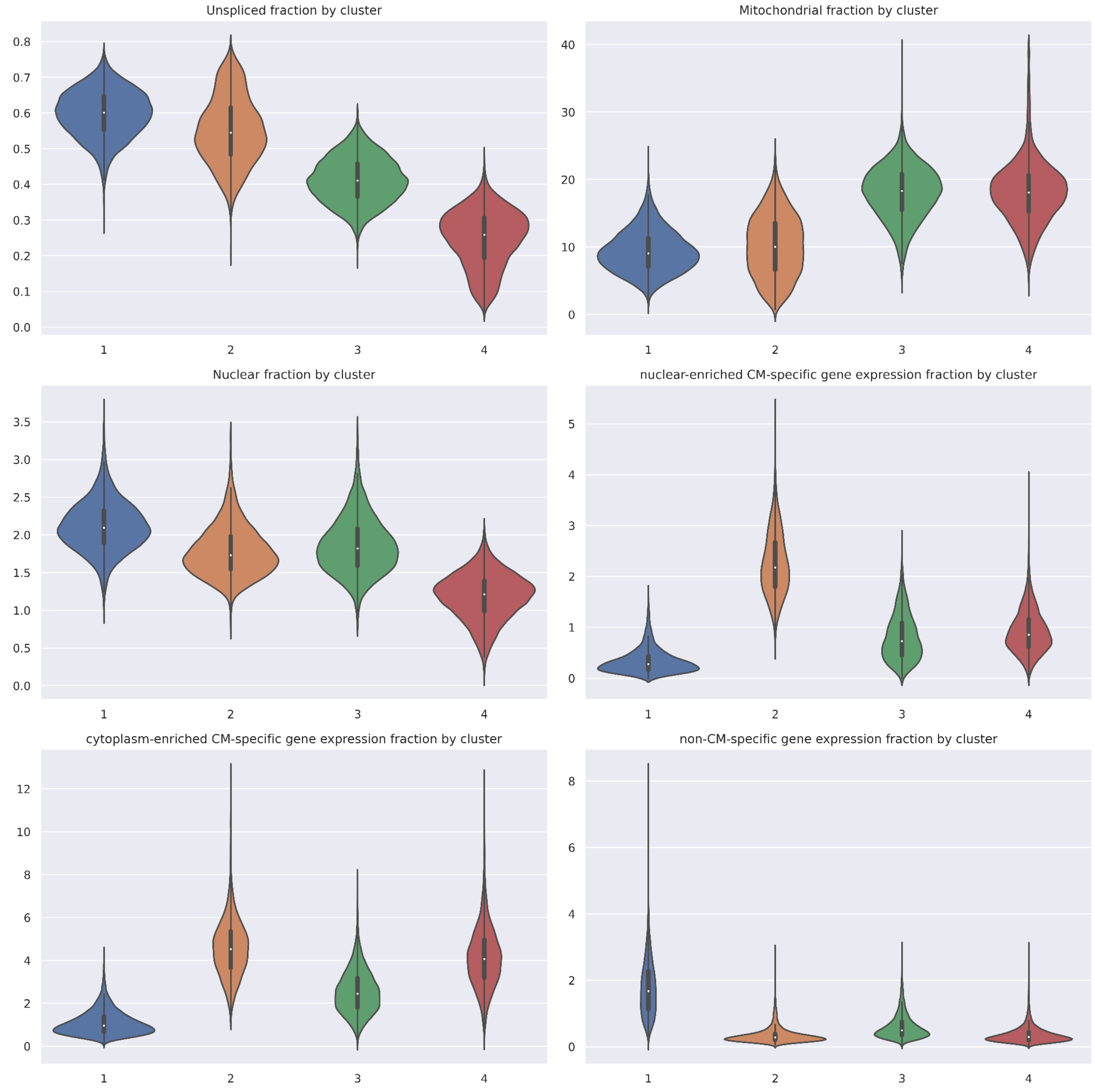


**Figure S5**. Distribution of clustering/quality metrics across k-means identified clusters.

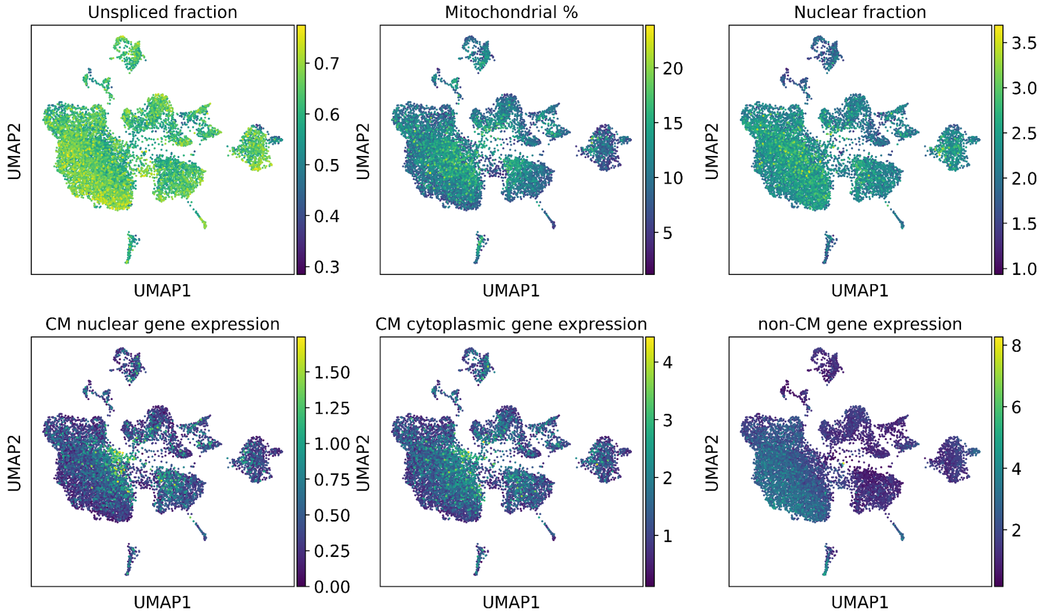


**Figure S6**. Distribution of clustering/quality metrics across UMAP of cluster 1, which corresponds to non-CM cell type populations.

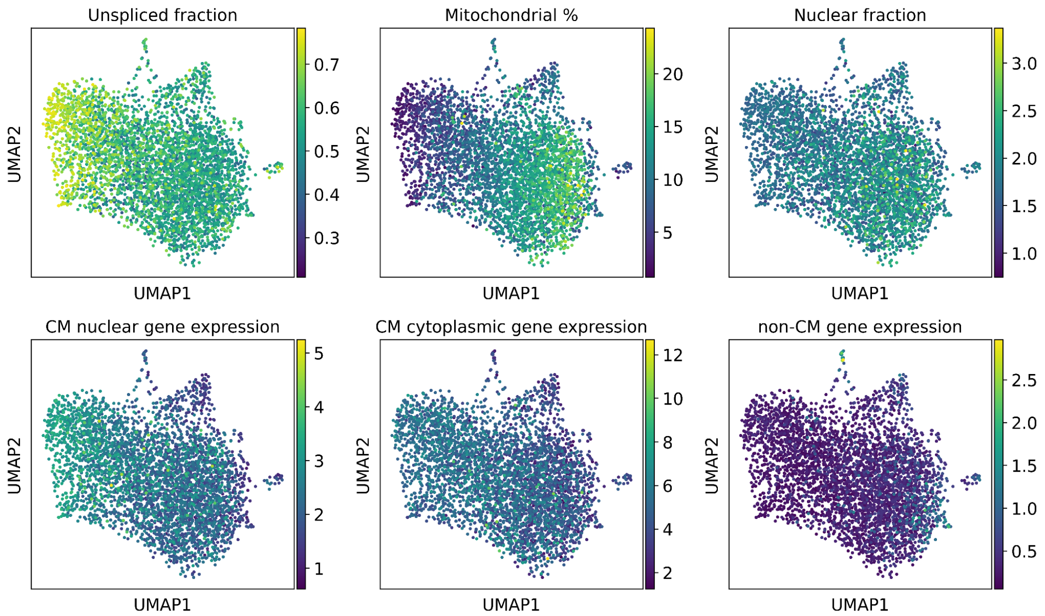


**Figure S7**. Distribution of clustering/quality metrics across UMAP of cluster 2, which corresponds to cardiomyocytes.

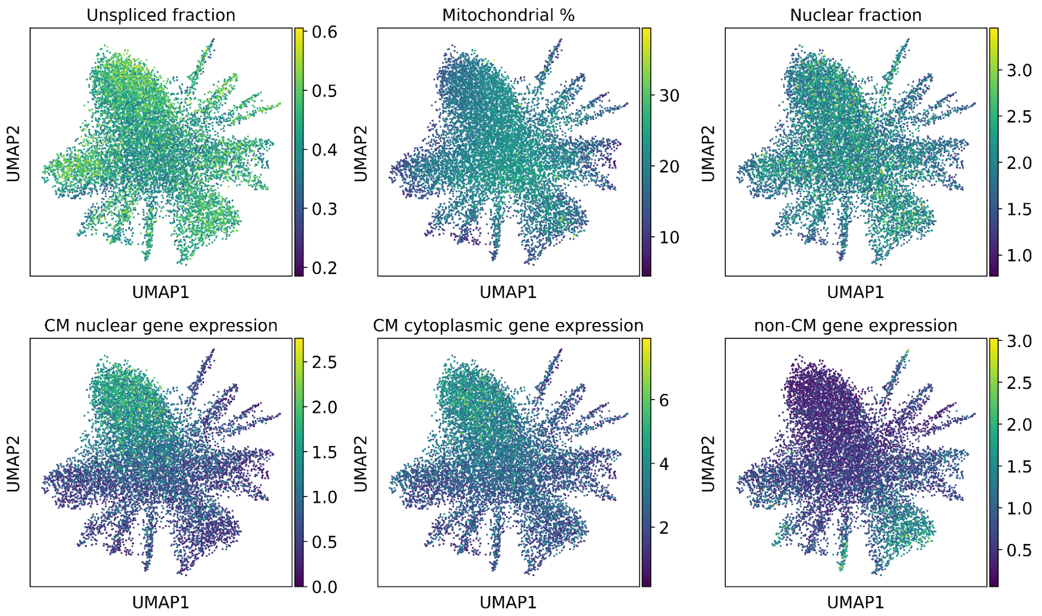


**Figure S8**. Distribution of clustering/quality metrics across UMAP of cluster 3, which corresponds to highly contaminated droplets.

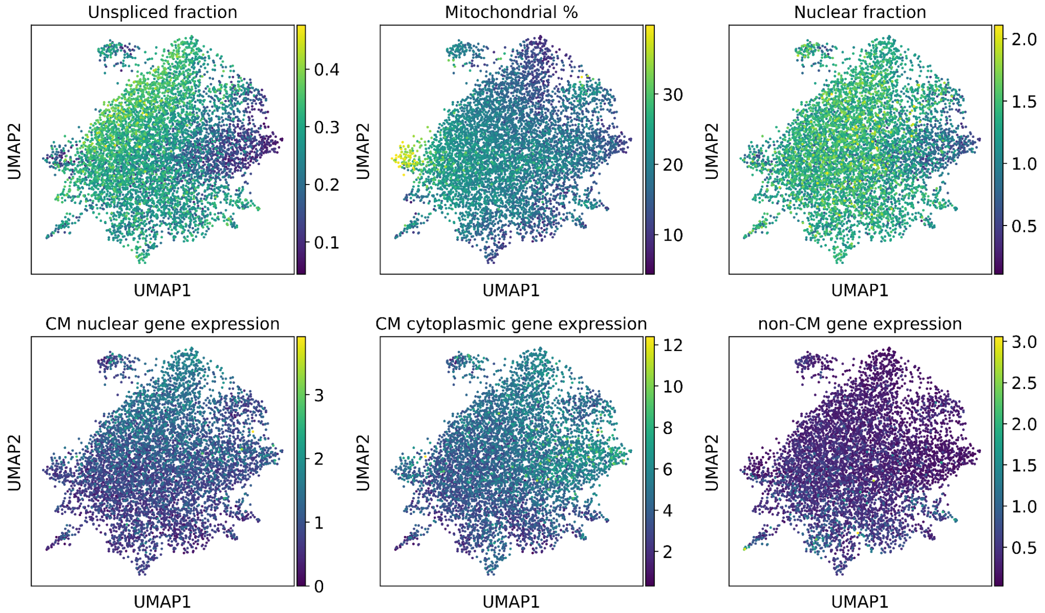


**Figure S9**. Distribution of clustering/quality metrics across UMAP of cluster 4, which corresponds to empty droplets.

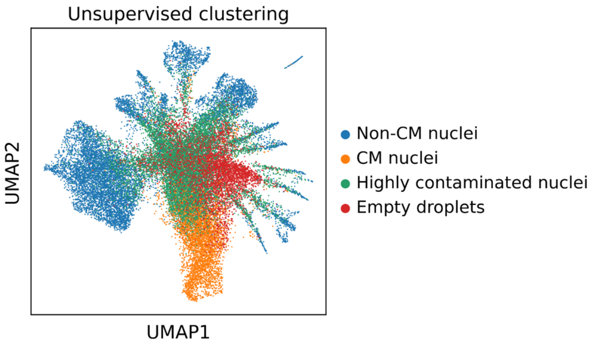


**Figure. S10**. Identified clusters on UMAP of sample CB-S00 from our previous study (doi: https://doi.org/10.1101/2021.06.23.449672).

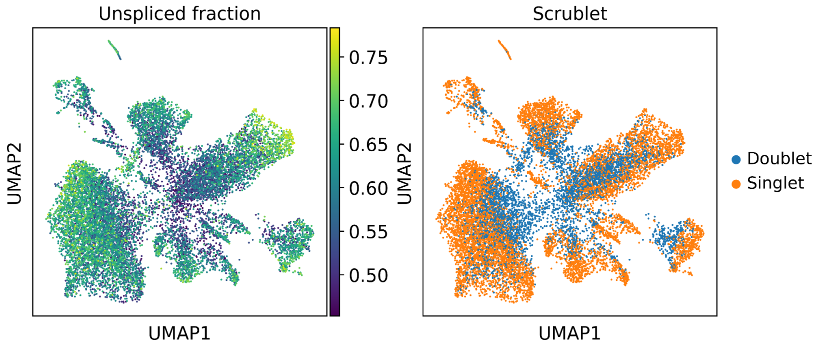


**Figure. S11**. Identified doublets display lower unspliced fraction than non-doublets.

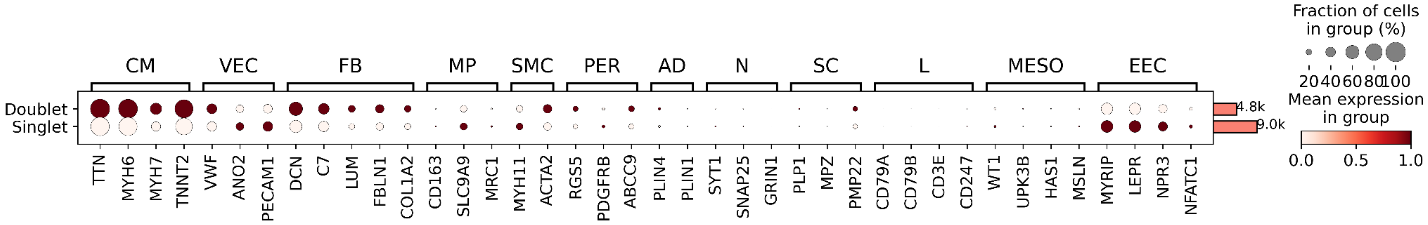


**Figure. S12**. Expression of marker genes in droplets identified as doublets show high expression of CM marker genes

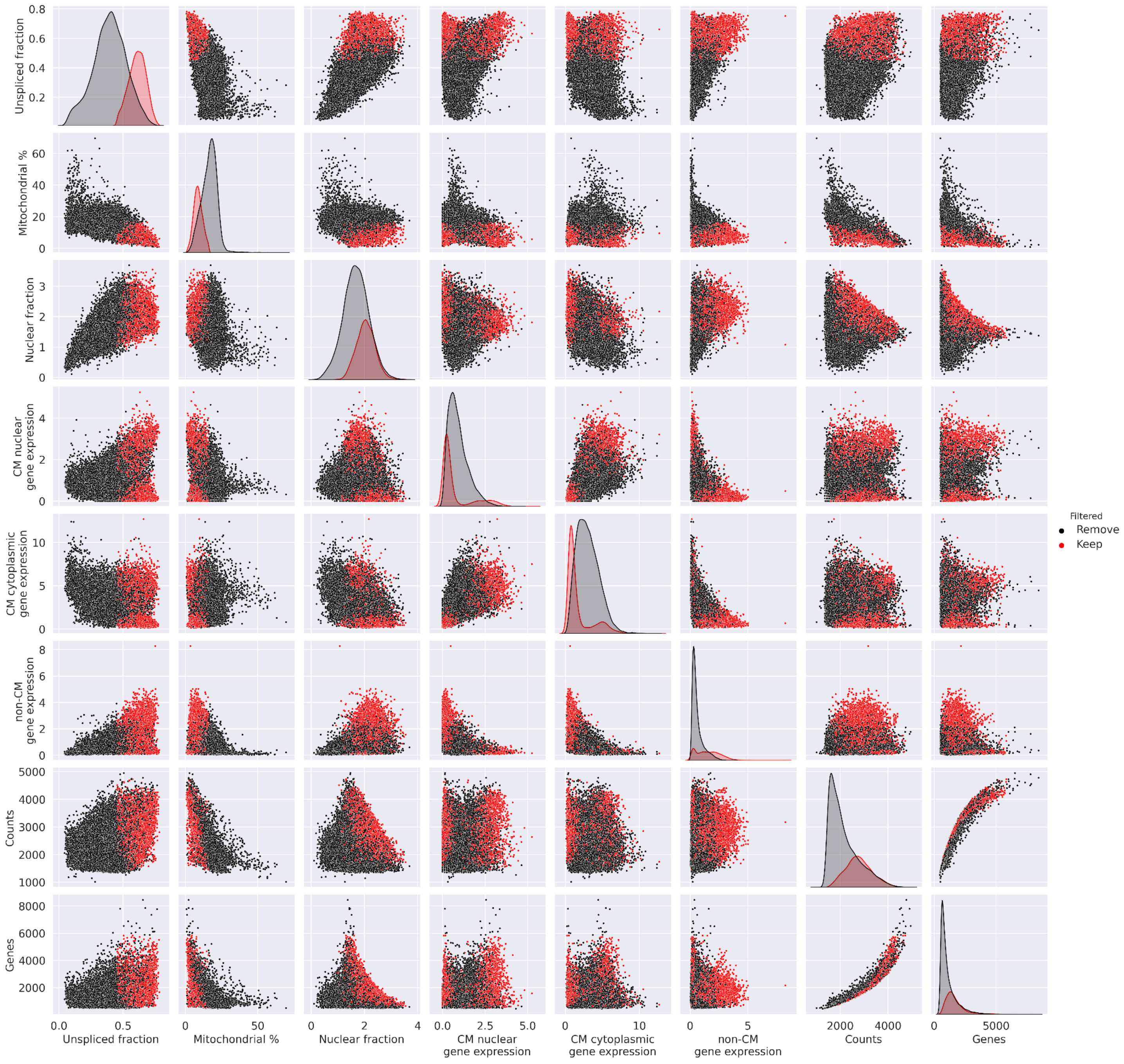


**Figure. S13**. Pair plot of quality metrics for kept and removed droplets. Every metric shows overlapping distributions of kept and removed nuclei, illustrating that no one metric alone could be used to remove the selection.

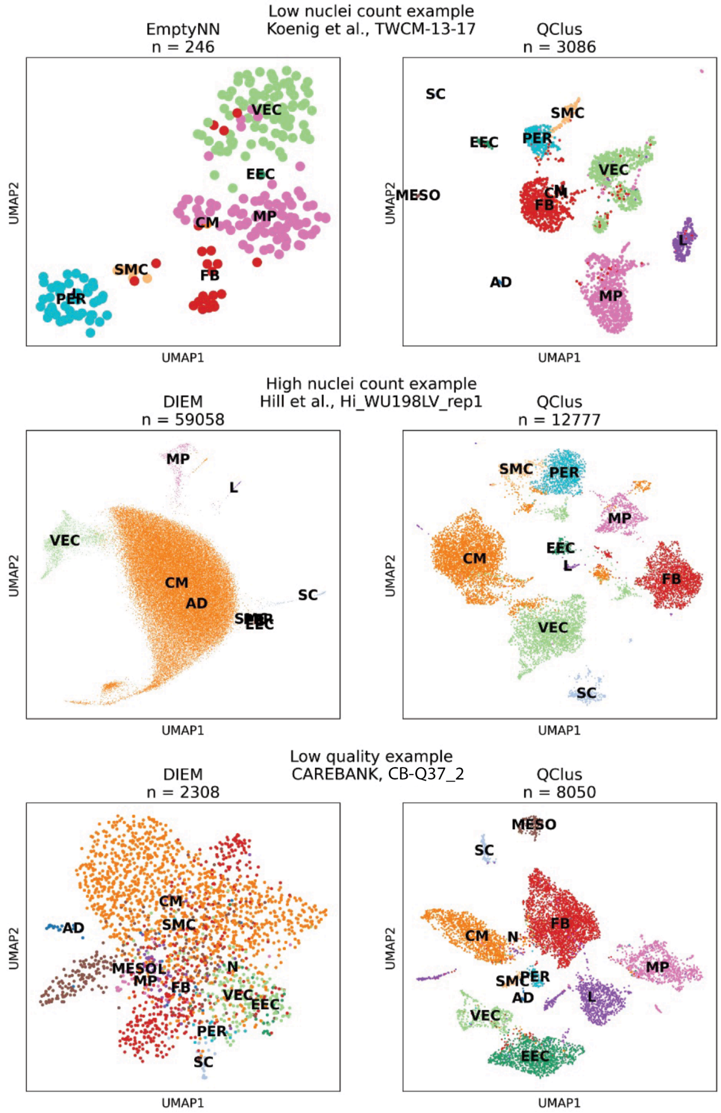


**Figure. S14**. Examples of processing failures (left column) against successful processing with QClus (right column) for three example samples.
